# Double asymmetric percolation drives a quadruple transition in sexual contact networks

**DOI:** 10.1101/784587

**Authors:** Hongyu Zheng, Xiangrui Zeng

## Abstract

Since 2007, ZIKV outbreaks have been occurring around the world. While ZIKV is mainly spread by mosquito vectors, transmission via sex activities enables the virus to spread in regions without mosquito vectors. Modeling the patterns of ZIKV outbreak in these regions remain challenging. We consider age as an asymmetric factor in transmitting ZIKV, in addition to gender as seen in previous literature, and modify the graph structure for better modeling of such patterns. We derived our results by both solving the underlying differential equations and simulation on population graph. Based on a double asymmetric percolation process on sexual contact networks. we discovered a quadruple ZIKV epidemic transition. Moreover, we explored the double asymmetric percolation on scale-free networks. Our work provides more insight into the ZIKV transmission dynamics through sexual contact networks, which may potentially provide better public health control and prevention means in a ZIKV outbreak.

## 1 Introduction

Zika virus (ZIKV) is a Flavivirus closely related to dengue. People infected with ZIKV may develop symptoms including fever, rash, and joint pain. Though the symptoms associated with ZIKV are generally mild, neurological complications have also been reported [10]. After its first identification in 1947, there were only a few human infections reported over a 60-year period. However, since 2007, ZIKV outbreaks have been reported in Yap island in many regions including Philippines, Pacific Area, and Americas [11, 9, 8, 21].

ZIKV is primarily transmitted through mosquito-borne infection. However, transmission via sex has also been reported [14]. This unique transmission dynamic requires specific biomedical research and mathematical modeling. In addition, the risks of transmission are suggested to be different with respect to gender and age [11].

Mathematical modeling has become an important tool in designing control and prevention means for infectious diseases [16]. A number of mathematical models of ZIKV transmission have been proposed. [15] developed a deterministic model of ZIKV transmission that accounts for both mosquito and sexual contact modes. [17] proposed a susceptible-exposed-infectious-removed (SEIR) framework. [5] proposed a model for identifying the most susceptible regions of ZIKV infection.

In this paper, we are interested in an asymmetric percolation model which accounts for the asymmetric duration of infectiousness between males and females [1]. Males can be infectious for over 180 days whereas females are infectious only for about 20 days [22, 24]. Therefore, the probability of ZIKV transmission is said to be asymmetric with respect to gender. Based on this observation, they modeled the ZIKV sexual transmission through the asymmetric percolation process on random sexual contact networks. By exactly solving their networks, they demonstrated a double transition, which identified two distinguishable thresholds for ZIKV to be endemic just based on sexual contact. The details of their model are shown in Section 2. We note here that their model and our proposed model only consider the sex transmission of ZIKV. This assumption can be applied to cases in which the infected mosquitoes are absent in the specific region but ZIKV is brought by foreign travelers who have sexual contacts with locals.

We modify and extend the existing model by incorporating more observations: 1) The risk of transmission is age-dependent. The ZIKV outbreak in Yap Island, Philippine, showed that for adults, increasing in age increases the susceptibility to ZIKV infection. Adults over 50 years old are nearly four times more likely to be infected and show ZIKV related symptoms than adults under 50 years old [11]. Therefore, besides gender, the age of individuals in the sexual contact network also causes an asymmetric probability of ZIKV transmission. We termed our new model as a double asymmetric percolation process on sexual contact networks. 2) Human sexual contact networks are shown to be scale-free [18, 13]. A scale-free network is one type of small-world networks whose degree distribution follows a power law. Scale-free networks have clustering coefficients much higher than random networks [3]. Survey has shown that the number of sexual partners decays as a scale-free power law [18]. Therefore, human sexual contact networks exhibit clustering property as scale-free networks. We investigated whether the multiple epidemic transitions (more than one distinguished epidemic thresholds) continued to exist when the underlying sexual contact networks were scale-free rather than random.

## 2 Related work

In this section, we will briefly describe the model present in *Asymmetric percolation drives a double transition in sexual contact networks* [1], the basis for our work. The authors present an asymmetric percolation model of ZIKV sexual transmission based on gender and sexual orientation.

First, they use nodes to represent individuals. Each node belongs to one of the six types (female/male and homo-/bi-/heterosexual), and is assigned to a number of sexual contacts. The proportion of the six types *w*_*i*_ is estimated with census and population research data. The contacts in the network are designed to be random. That is, for a node, the number of sexual contacts, *k*, follows a Poisson distribution with < *k* >= 5, regardless of its type [4].

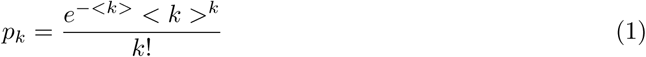

Second, edges are created randomly between pairs of nodes constrained by their types. For example, a homosexual female can have sexual contacts with homosexual females as well as bisexual females. Based on the proportion of node types *ω*_*i*_, *α*_*j*|*i*_, the probability that a neighbor of a node of type *i* is of type *j*, is calculated from *ω*_*i*_ based on the principle that the proportion of outgoing edges to a certain type should loosely match its population size and the resulting distribution favors homosexual people over bisexual peoples. For this report, we purpose no change to the values of *α* for comparison purposes, but better mechanisms for deciding *αj*|*i* might be possible. Due to the nature of the half-edge matching algorithm, it is required that *ω*_*i*_ × *α*_*j*|*i*_ = *ω*_*j*_ × *α*_*i*|*j*_ for all pairs (*i, j*) ∈ [*N*]^2^, where *N* denotes the set of six possible types.

Then, the transmissibility, *T*_*ij*_ is defined to be the probability of transmission from a node of type *i* to a node of type *j*. Due to the fact that males have much longer ZIKV infectiousness period, the transmissibility from a male to his sexual partner is higher than from a female to her sexual partner. Therefore, they introduced an asymmetry parameter *a* to downscale the transmissibility when *i* is female. Specifically,

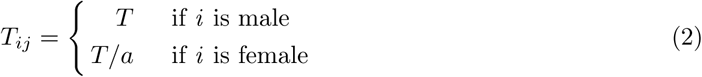

Based on the transmissibility *T*_*ij*_ and the simulated network, the epidemic threshold and the expected final size of outbreaks can be computed [2] as follows:

In a random population graph, a component of population graph is called a giant component if it contains a constant fraction of the entire graph (whole population), and is called a small component otherwise [23]. In a hypothetical scenario, one person of a certain type, denoted *i*, is first infected, then with a probability of *T*_*ij*_ the disease spreads to its neighbors of type *j*. The process continues from the newly infected person until no more individuals fall victim. The set of infected individuals is then a component of the population graph, and the event is called an outbreak if the component is a giant one.

Three metrics of interest are proposed, all defined on single classes and on the population as a whole, and take the composition of graph and choice of nodes as underlying random space:

- *P*_*i*_: Probability of a person of type *i* starting an outbreak.
- *S*_*i*_: Expected fraction of type *i* population involved in an outbreak.
- < *s* >_*i*_: Expected number of victims of type *i* in a non-outbreak.

## 3 Methods

### 3.1 New Models

#### Double Asymmetric Model

To incorporate the observation of the age dependency of ZIKV transmissibility, we further classified the nodes in a sexually active population network into more types. There are now 12 types of nodes defined by age, gender, sexual orientation (old/young, female/male, homo-/bi-/heterosexual). The proportion of each type of nodes, *w*_*i*_, is estimated by surveys of ageing [19] and summarized in Table 3.1. An individual is defined as old when he or she is over 50 years old.

**Table 1:** Description of node types *i* and their relative proportion *w*_*i*_ in sexually active population. These values are used in the numerical simulation.

|  |  |  |  |  |
| --- | --- | --- | --- | --- |
| Description<br>$i$<br>$w_i$ | Old Homo. males<br>0<br>0.625% | Old Homo. females<br>1<br>0.625% | Old Bi. males<br>2<br>0.375% | Old Bi. females<br>3<br>0.375% |
| Description<br>$i$<br>$w_i$ | Old Hetero. males<br>4<br>11.5% | Old Hetero. females<br>5<br>11.5% | Young Homo. males<br>6<br>1.875% | Young Homo. females<br>7<br>1.875% |
| Description<br>$i$<br>$w_i$ | Young Bi. males<br>8<br>1.125% | Young Bi. females<br>9<br>1.125% | Young Hetero. males<br>10<br>34.5% | Young Hetero. females<br>11<br>34.5% |

**Table 2:**
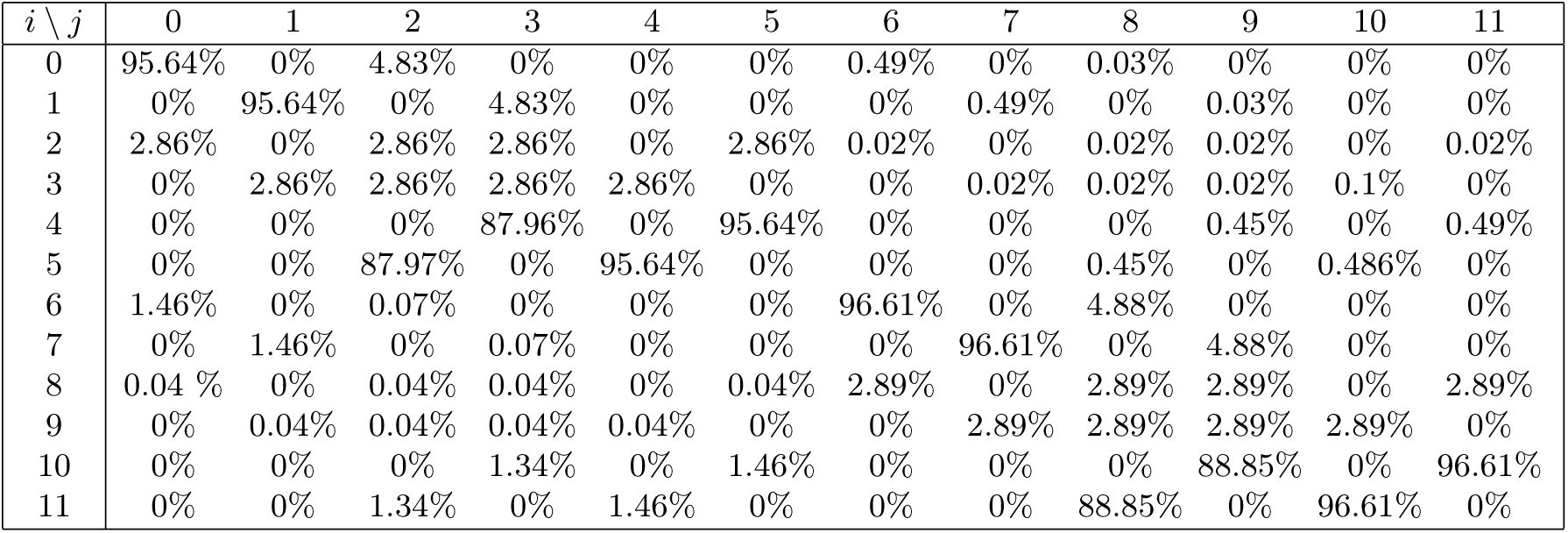
Value for *α*_*i*|*j*_ in new models. These values are generated based on affinity model proposed above and is used in all numerical simulations.

| $i \setminus j$ | 0 | 1 | 2 | 3 | 4 | 5 | 6 | 7 | 8 | 9 | 10 | 11 |
| --- | --- | --- | --- | --- | --- | --- | --- | --- | --- | --- | --- | --- |
| 0 | 95.64% | 0% | 4.83% | 0% | 0% | 0% | 0.49% | 0% | 0.03% | 0% | 0% | 0% |
| 1 | 0% | 95.64% | 0% | 4.83% | 0% | 0% | 0% | 0.49% | 0% | 0.03% | 0% | 0% |
| 2 | 2.86% | 0% | 2.86% | 2.86% | 0% | 2.86% | 0.02% | 0% | 0.02% | 0.02% | 0% | 0.02% |
| 3 | 0% | 2.86% | 2.86% | 2.86% | 2.86% | 0% | 0% | 0.02% | 0.02% | 0.02% | 0.1% | 0% |
| 4 | 0% | 0% | 0% | 87.96% | 0% | 95.64% | 0% | 0% | 0% | 0.45% | 0% | 0.49% |
| 5 | 0% | 0% | 87.97% | 0% | 95.64% | 0% | 0% | 0% | 0.45% | 0% | 0.486% | 0% |
| 6 | 1.46% | 0% | 0.07% | 0% | 0% | 0% | 96.61% | 0% | 4.88% | 0% | 0% | 0% |
| 7 | 0% | 1.46% | 0% | 0.07% | 0% | 0% | 0% | 96.61% | 0% | 4.88% | 0% | 0% |
| 8 | 0.04 % | 0% | 0.04% | 0.04% | 0% | 0.04% | 2.89% | 0% | 2.89% | 2.89% | 0% | 2.89% |
| 9 | 0% | 0.04% | 0.04% | 0.04% | 0.04% | 0% | 0% | 2.89% | 2.89% | 2.89% | 2.89% | 0% |
| 10 | 0% | 0% | 0% | 1.34% | 0% | 1.46% | 0% | 0% | 0% | 88.85% | 0% | 96.61% |
| 11 | 0% | 0% | 1.34% | 0% | 1.46% | 0% | 0% | 0% | 88.85% | 0% | 96.61% | 0% |

We also depart from the principle of *α*_*i*|*j*_ reflecting the proportion of population *ω*_*i*_ and introduce the idea of affinity, which means sexual interactions are much more likely to happen within an age group, than between the group [12]. This is reflected in the design of *α*_*i*|*j*_:

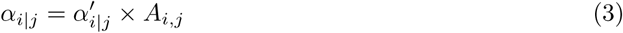

Where *α*′ represents the original probability without age group, and *A*_*i,j*_ is an affinity factor that is chosen to seperate the two groups

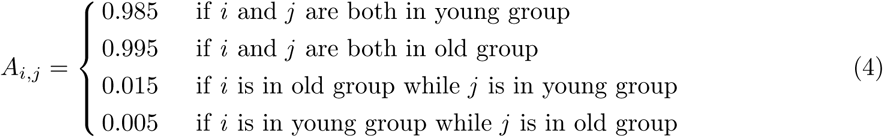

while keeping *ω*_*i*_ × *α*_*j*|*i*_ = *ω*_*j*_ × *α*_*i*|*j*_ as required by half-edge matching algorithm. The actual value used in this part of simulations can be found in Table 3.4. For example, *α*_1|3_ = 4.83% means that in our numerical simulation, on average, 4.83% of an old homosexual female’s sex partners are old bisexual females.

We introduce a new asymmetry parameter *b*. In particular, for every *i, j* ∈ *N*, we increase the chance of transmission *T*_*ij*_ by a factor *b* when *j* is old (over 50 years old). *b* is chosen to be smaller than *a* in our numerical simulation. The modified transmissibility is

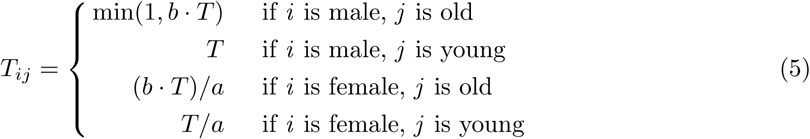

#### Alternative Degree Distributions

Moreover, we are also interested in other feasible models of sexual activities. The paper [13] discovers power-law degree distribution of sexual contact network, so we also generated scale-free networks to test the model. Instead of a Poisson distribution, the degree of each node is sampled from:

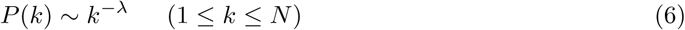

Where *k* ≥ 1 and *λ* is taken to be approximately 2.050 in the first experiments to match the Poisson model with E(*k*) ≈ 5 for *N* ≈ 10^4^, total population. We show examples of random networks versus scale-free networks in Figure 1. Scale-free networks exhibit more clustering properties resembling a human sexual contact network. This value is also consistent with the fact that most real-world networks have a *λ* factor between 2 and 3. In later experiments, we specifically consider the case where *λ* > 3 for it exhibits different behavior.

**Figure 1:**
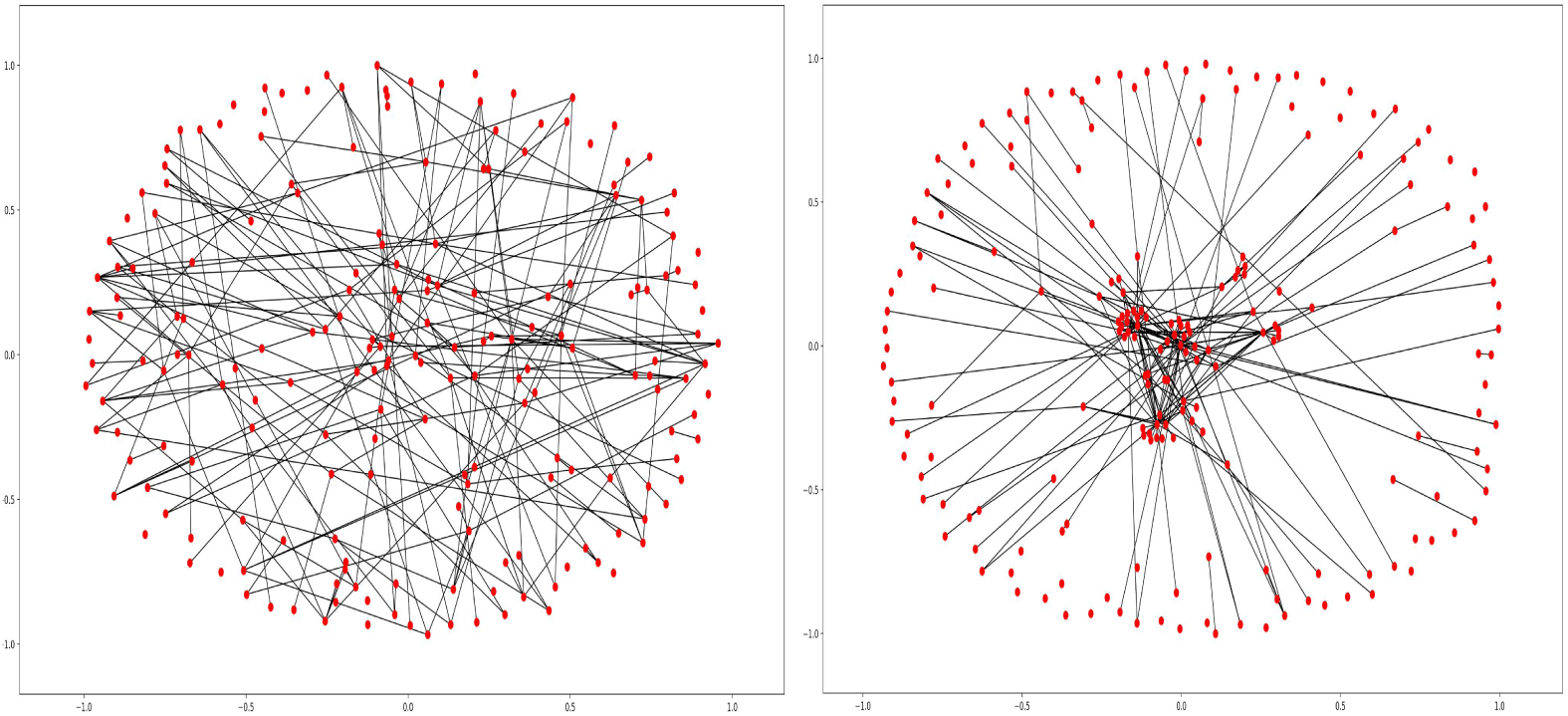
Examples of Left: a random network; Right: a scale-free network.

In the following sections, we give an overview of two different methods to solve the values of *P, S* and < *s* > under different values of *T*. The idea of turning the problem of solving *P* and *S* into solving for fixed points of a vector function **f**(**x**) shall be attributed to [2] for a more general formalism, and the elegant solution for solving < *s* > without complicated marginalization comes from [1]. The stab matching protocol comes from [2](See Implementation Details for more information) and is specified in [1], and the process of validating analytical solutions with simulation results is partly inspired by [1]. We also make changes to the various parts for better demonstration.

#### 3.2 Solution by Iteration

For all three metrics, analytical solutions are difficult to compute. However, an argument called “self-consistency” can be invoked for estimating the values. Here we demonstrate the technique on solving *P*_*i*_. Due to limitations on space, we only discuss the high-level ideas behind and skip actual calculations. We introduce several terms:

- A bold variable **x** is a vector of length *n*, number of node types(6 in the original model).
- *g*_*i*_(**x**) is the probability-generating function of composition of direct successors for a node of type *i*(a direct successor is a node can be visited by using one outgoing edge from current node). The probability that a node of type *i* have exactly *v*_*k*_ direct successor of type *k* is given by the coefficient before term 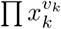 in *g*_*i*_(**x**). Normalization requires *g*_*i*_(**1**) = 1.
- *f*_*li*_(**x**) is the probability-generating function of composition of direct successors of a type *i* node that is reached from an edge of type *l*. This is analogous to the “excess degree” in classical percolation theory. However, since all *f*_*li*_ are equal for same *i* thanks to the simple structure, we drop the first subscript and simply use *f*_*i*_.

*f*_*i*_(**x**) and *g*_*i*_(**x**) have a closed form and can be calculated efficiently when **x** is given. For sake of completeness, *f* and *g* are defined as

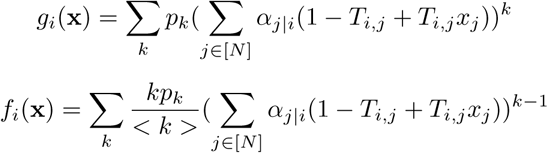

where Σ_*k*_ iterates over the possible values degree distribution, *p*_*k*_ denotes probability of a degree *k* node, and < *k* >= Σ_*k*_(*kp*_*k*_), average degree of the graph.

Next, we define *u*_*i*_ to be the probability of a type *i* node visited from some edge **NOT** starting an outbreak(not in a giant component).

The key assumption here is that the graph is “locally tree-like”, meaning that there will be no loops in the random network. Given this, the probability of this type *i* node **NOT** starting an outbreak is exactly the probability that any of its direct successors does not satisfy the property, independent of each other(and the probability for a direct successor of type *j* not satisfying the property is given by exactly *u*_*j*_). Iterating over all possible configuration of successors of the node, it leads to the conclusion 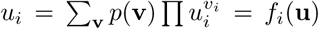, so **u** is a fixed point of function **f**(**x**) = [*f*_*i*_(**x**)].

*P*_*i*_, the probability of a person of type *i* starting an outbreak(note the difference with *u*_*i*_ on how the node is picked, thus the different underlying random space) is given by *P*_*i*_ = 1 − *g*_*i*_(**u**), following similar argument over all possible configuration of direct successors of a type *i* node, which is now generated by *g*_*i*_(**x**).

*S*_*i*_ can be derived in a similar iterative manner. The derivation of < *s* >_*i*_ is more involved and we shall redirect the reader to the original paper for details.

### 3.3 Solution by Simulation

For a realistic estimate, we first generate a population graph of size 10^4^ with predefined degree distribution, population composition, and transmissibility between node types. The graph is then condensed, that is, we partition the graph into strongly connected components (SCC) and contract each SCC to a single node. This greatly reduces the computational cost, as we can treat each SCC as a whole when simulating infections. An outbreak is defined as an infection over 1 percent of the population. The condensed graph is now a Directed Acyclic Graph (DAG) and we calculate the set of successors (number of infections if starting from the current node) for each condensed node.

We calculate the three main metrics as follows.

- *P*_*i*_ is calculated as the average proportion of type *i* nodes in a outbreak.
- *S*_*i*_ is calculated exactly the same way as *P*_*i*_, except all edges have their direction reversed.
- < *s* >_*i*_ is calculated as the average number of type *i* nodes in a non-outbreak.

Later, we shall show this definition makes sense and matches theoretical predictions.

### 3.4 Implementation details

#### Simulation

To build the population graph, we utilize the DiGraph class in NetworkX package as the underlying structure, and employ the “stab-matching” procedure to generate a graph with asymptotically correct degree distribution and affinities. First, the desired degree for each node is drawn from input distribution. For each node *x* with desired degree *d*, we add *d* half-edges labeled with *x* to a list. With probability *w*_*i*_*α*_*j*|*i*_, we pick two half-edges with type *i* and *j*, and establish a link between the nodes. This process is repeated until there are no more feasible half-edges to match.

Then, to simulate transmission, we break each edge generated above into two directed edges. A edge from *i* to *j* is discarded with probability 1 − *T*_*ab*_ where *a* and *b* are the types of *i* and *j*. Due to technical limitations on iterating over incoming edges, we build two graphs, one with the aforementioned method, and a second one that is identical to the first except all edge reversed. Depth First Search(DFS) was used to iterate over all successors(victims) from a particular node and calculate *P*. To calculate *S* and < *s* >, the reversed graph is used.

#### Parallelization

In addition to Section 3.1, The final result is averaged over 100 − 500 instantiations of the population graph to count for intrinsic randomness of the model. Since the calculation requires we knowing the exact composition of each subgraph generated from every people, a full parameter scan(on *T*) can take up to hours. We use a cluster to run multiple instances simultaneously(up to 100), each with its own generated population graph, and use another program to collect the results.

The simulator is implemented in Python 3. As a side note, we have contacted the authors of the original paper [1] and confirmed our simulation method.

#### Solution by Iteration

We also use self-consistency argument to transform the simulation into finding fixed points for vector functions. Changes are made to accommodate for changes in a population model, while the high-level idea is similar and the self-consistency argument remains the core.

In practice, we start with *x*_*i*_ = 0.5 for all dimensions, and to find a fixed point we iterate **x**_*t*+1_ = **f**(**x**_*t*_) until converge, defined by |**x**_*t*+1_ − **x**_*t*_|_∞_ < *ϵ. f* and *g* are polynomial functions of **x** when the network structure is fixed, so this is effectively a Fixed Point Iteration methods for finding a root.

Since all terms are normalized and all positive, *x*_*i*_ = 1 for all dimensions will always be a fixed point(*u*_*i*_ = 1 indicates no point is in the giant component, thus there is no giant component). Furthermore, for *T* → 0 it’s the only solution and a stable one. However, as *T* increases, a bifurcation occurs and a second solution where some *x*_*i*_ < 1 will show up. This indicates the emergence of a giant component. After the bifurcation occur, *x*_*i*_ = 1 is also no longer a stable fixed point(due to the fact that *f* is convex over [0, 1]^*N*^). This leads to the conclusion that the algorithm is guaranteed to return the giant component or return the *x*_*i*_ = 1 vector if there is no giant component.

We take *ϵ* = 10^−8^ in all cases. The solver is implemented in Python 3, and finishes a parameter scanning on *T* within 10 minutes on most instances.

#### Visualization

We designed a simple format to store results from both iteration and simulations. The result file is fed to another program which generates the curve of *P, S* and < *s* > for each value of *T*, averaged if multiple experiments are done with the same value. The resulting graph is exported as PNG image file and used in this report. For each image, approximately 100 data points was used. We perform no smoothing or averaging between different *T* s.

#### Code Availability

All codes are available upon request, including code for model construction, both solutions, visualizer, and script required to dispatch tasks on the cluster.

#### Calculated Values of Transitions

To avoid cluttering, Here we list the complete table of *α*_*i*|*j*_ used in the double asymmetric model.

## 4 Results

### 4.1 Reproducing original results

In their analytical calculation, two epidemic threshold can be clearly seen in Figure 2. They demonstrated that due to the asymmetric ZIKV transmissibility, there will be two outbreak epidemic thresholds, a smaller one for the homosexual male population and a larger one for other individuals in the network.

**Figure 2:**
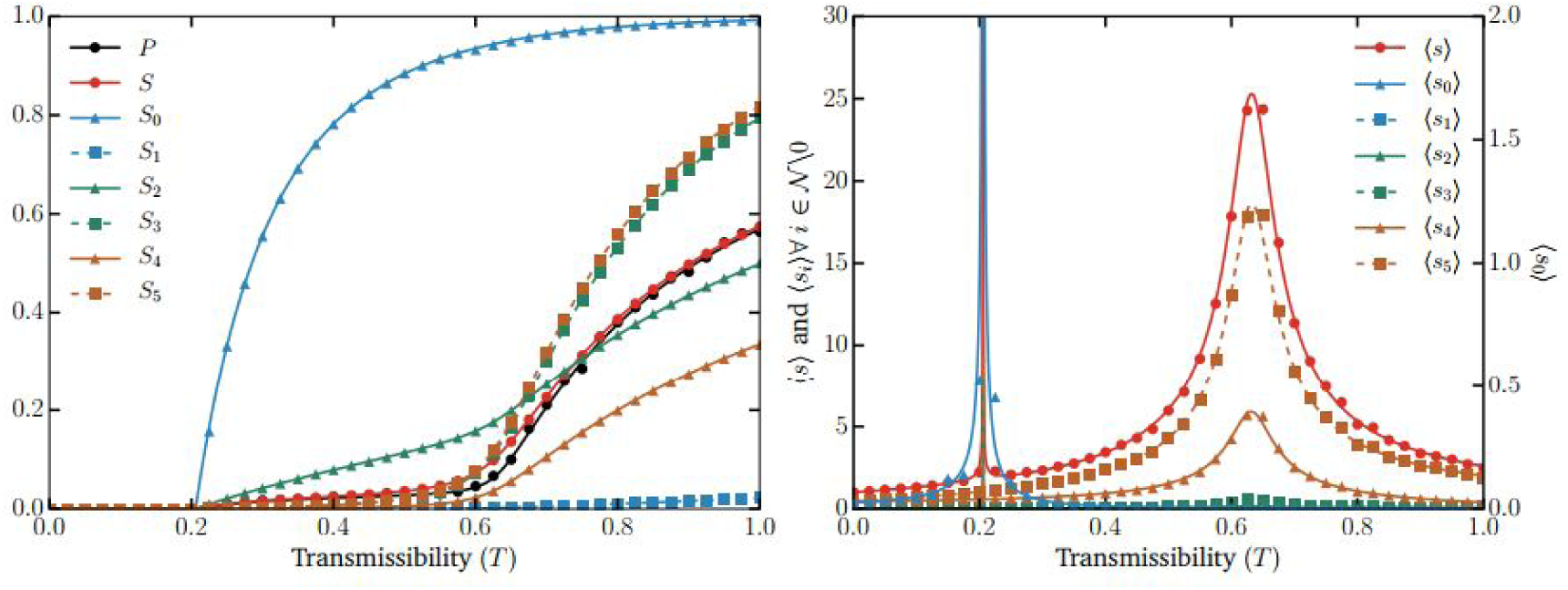
Analytical results of Left: The final relative size of outbreak by 6 types of nodes; Right: Two epidemic thresholds represented by <s>. Figure from [1] and reproduced below.

We implemented the analytical calculation and obtained similar results to their original analytical results. Our result is shown in Figure 3.

**Figure 3:**
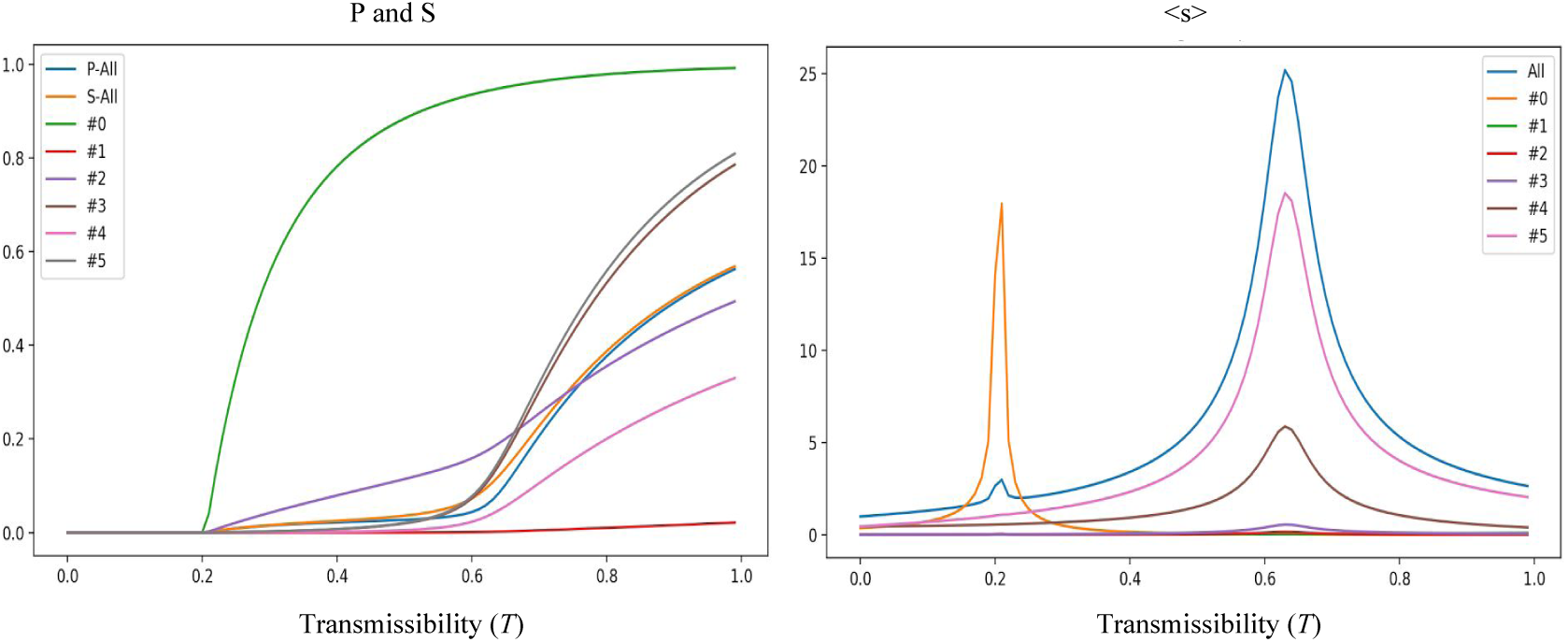
Reproduction of results in Figure 2. < *s*_0_ > is scaled up 15x to show the transition.

### 4.2 Double asymmetric percolation model results

Our proposed double asymmetric percolation model takes into consideration the double asymmetric ZIKV transmissbility concerning both gender and age. The original six types of nodes are subdivided into twelve types of new nodes.

For the new model, we plotted the three metrics of interest in Figure 4. Based on the overall average number of nodes in non-outbreaks (< *s* >), the blue line in Figure 4 right, we can clearly see three peaks, one around *T* = 0.05, one around *T* = 0.15, and one around *T* = 0.625. Therefore, there exist at least three global epidemic thresholds.

**Figure 4:**
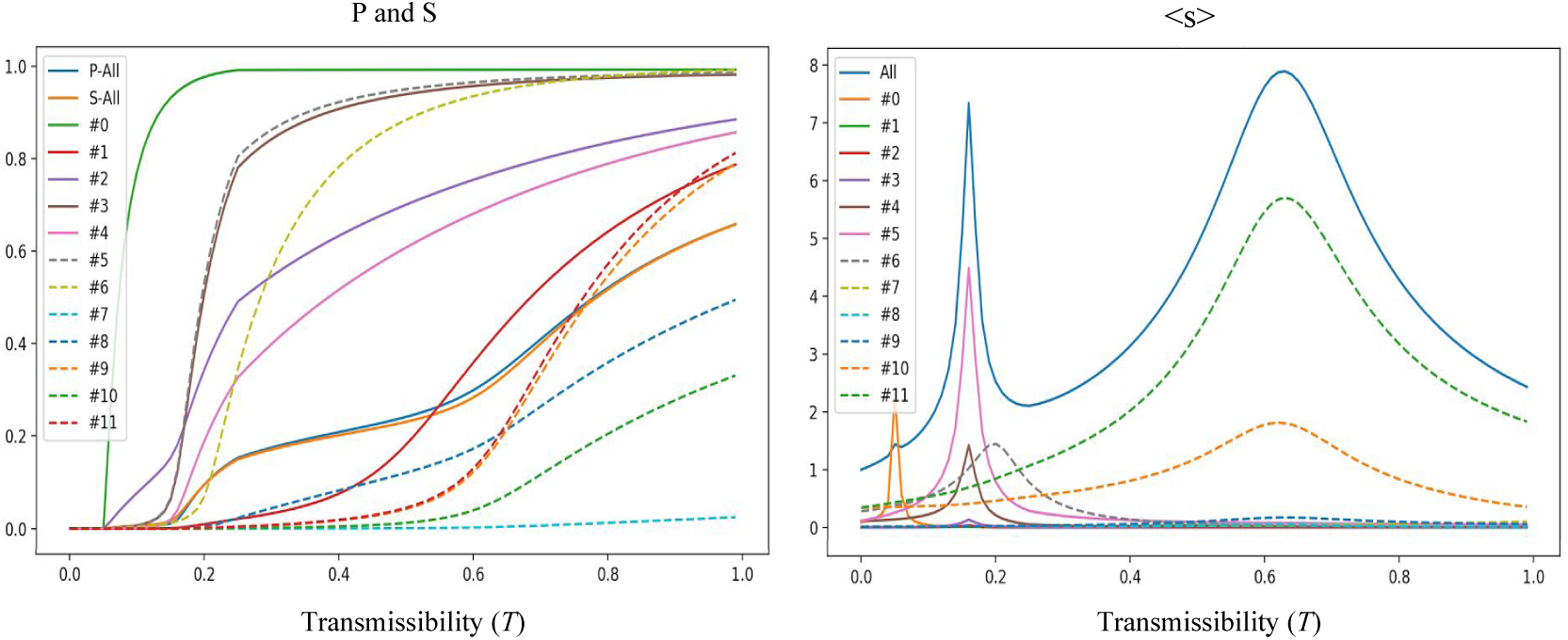
Analytical results of new model Left: The final relative size of outbreak by 12 types of nodes; Right: Epidemic thresholds represented by < *s* > and < *s*_*i*_ >.Be noted that < *s*_0_ > and < *s*_6_ > is scaled up 15x. This applies to all images below.

By analyzing the components of < *s* >, we found four distinguished peaks, one for old homo-sexual males at around *T* = 0.05, one mainly for old heterosexual males and females at around *T* = 0.15, one for young homosexual males at around *T* = 0.2, and one for young heterosexual males and females as around *T* = 0.625.

Furthermore, we validated our results using simulation on realistic data. The details of our simulation procedure can be found in Section 3.3. We simulated a population of 10^4^ individuals. All parameters used in simulation, *a, b, w*_*i*_, and *α*_*ij*_ are consistent with what we used in the analytical solution. Figure 5 shows the results of simulation study. Four peaks similar to analytical results in Figure 4 right, were obtained.(Note that colors are different.)

**Figure 5:**
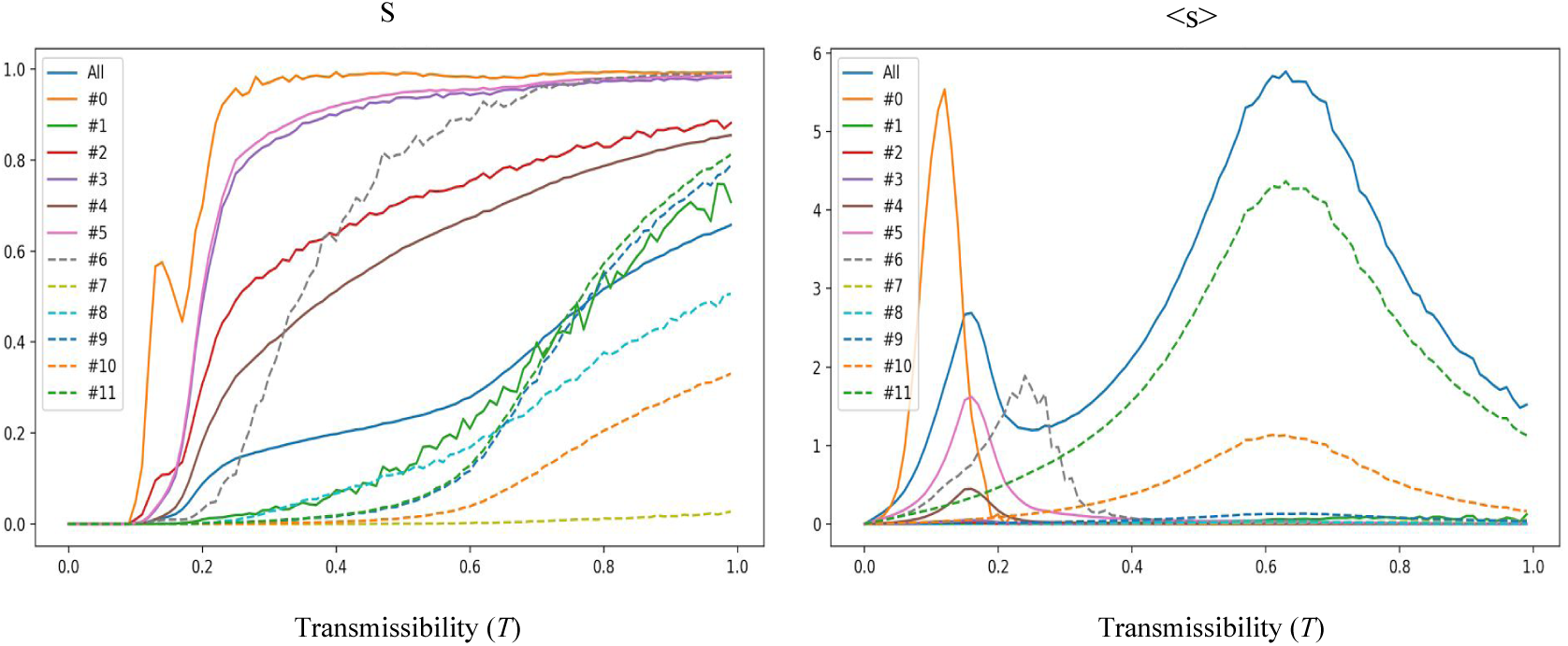
Simulation results of new model Left: The final relative size of outbreak by 12 types of nodes; Right: Epidemic thresholds represented by < *s* > and < *s*_*i*_ >.

#### Comparing Simulation results with Analytical Solutions

We produce highly similar trajectories for *S* and *P* with expected fluctuations around theoretical value in this case, validating our algorithm.

However, one can notice that while the position of peaks is mostly correct in the trajectories of < *s* >, the width and amplitude is off. This does not signal that our simulation is wrong; It’s well expected with this size of the population. Due to limited computational power, we decided that *n* ≈ 10^4^ is a good trade-off between the number of data points accessible from simulation (speed of simulation) and a larger population (in order to have a better separation of giant components and small components), both serving the ultimate target of higher accuracy.

To provide a bit more insight, in our current implementation, it’s almost impossible to judge if a component with 100 pops is a giant one or a small one. The emergence of small components is largely independent of graph size as can be seen in the derivations, but the large components are fractions of the total population. As a rule of thumb, we set the cutoff to 1 percent of the total population, which means that if *n* > 10^4^, the component will be classified as a giant one, and if *n* < 10^4^ it’ll be a small one. However, close resemblance in other parts and in general should prove the validity of the simulation approach, and we are confident that the problems shall diminish as we have access to more computational power and better algorithms for the simulation.

### 4.3 Double asymmetric percolation on scale-free networks

We further investigate whether the multiple epidemic thresholds continue to exist in scale-free networks.

In this part, since we are investigating the existence of a transition, we will not use simulations and depend on analytical solutions. The reason behind is that we are investigating whether an epidemic threshold exists in such cases, and in simulations, a component with less than 1 percent population is always classified as a small component(even if it should be a giant one by definition), which means that a “transition” will always present in simulation due to the component growing beyond 1 percent threshold, and it would be the best to rule out such artifacts. On the other hand, power-law distribution has a much longer tail than Poisson distribution, which means observations over simulations will converge much slower.

After modifying the degree distribution of all nodes to be scale-free, we plotted the three metrics of interest in Figure 6. The scale-free parameter *λ* is set to be 2.050 in this computation.

**Figure 6:**
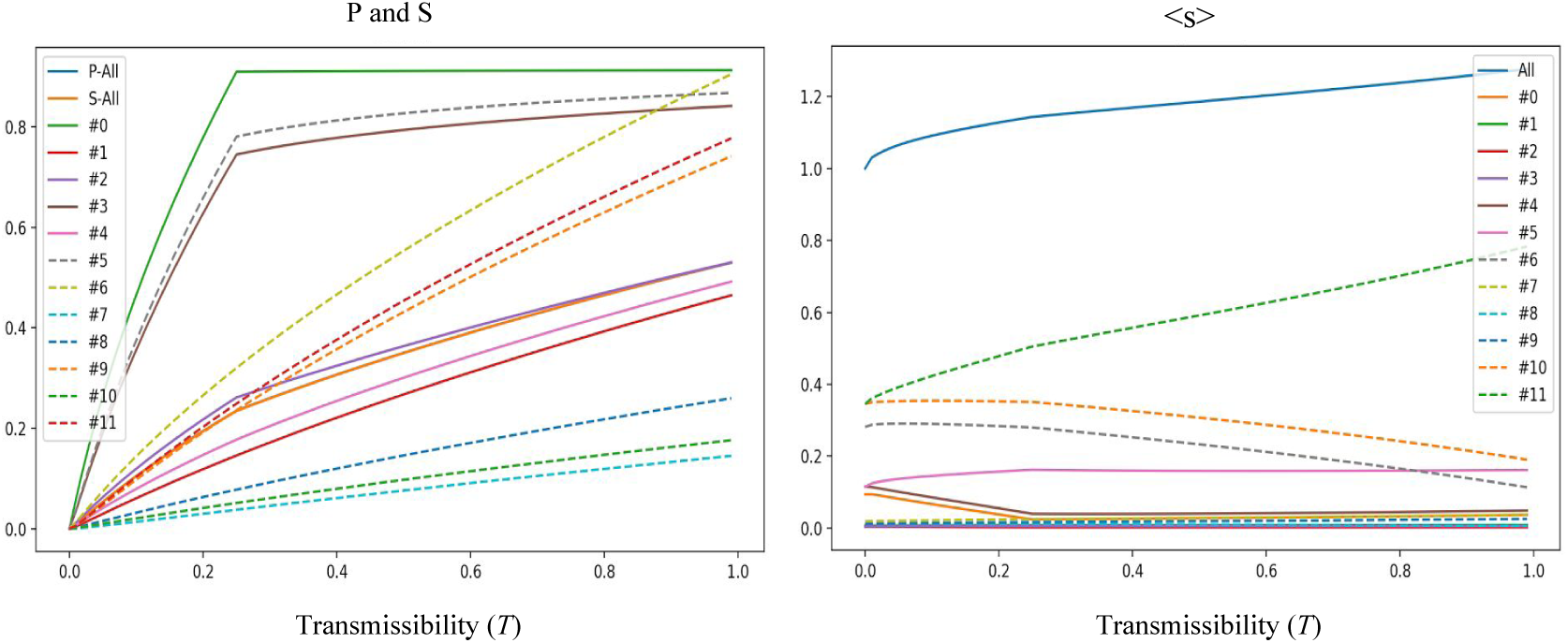
Analytical results of new model on scale-free networks *λ* = 2.050 Left: The final relative size of outbreak by 12 types of nodes; Right: Epidemic thresholds represented by < *s* > and < *s*_*i*_ >.

Figure 6 showed that on scale-free networks, all the epidemic thresholds (peaks in <s>) vanished. We note here that some of the elbow points occurring at *T* = 0.25 are due to the definition of *T*_*ij*_ being min(1, *b · T*), if *i* is male and *j* is old. When *b* = 4, *T*_*ij*_ = 1 for all *T* ≥ 0.25.

To further investigate this interesting phenomena, we tuned the value of the scale-free parameter *λ*. We found that when *λ* is greater than 3, peaks reappeared in the < *s* > graph.

We plotted the three metrics of interest in Figure 7. The scale-free parameter *λ* is set to be 3.1 in this computation.

**Figure 7:**
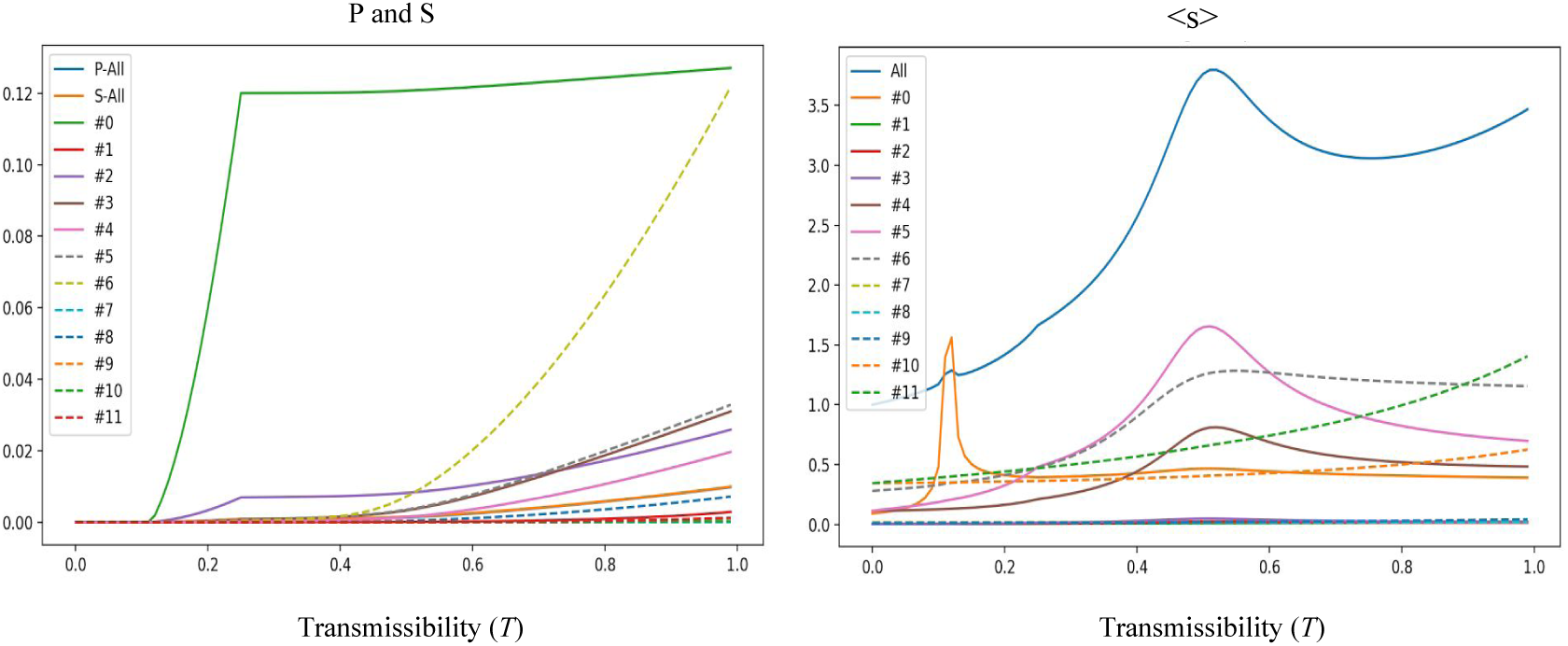
Analytical results of new model on scale-free networks *λ* = 3.1 Left: The final relative size of outbreak by 12 types of nodes; Right: Epidemic thresholds represented by < *s* > and < *s*_*i*_ >.

A further discussion of this interesting phenomena can be found in the Discussion section.

## 5 Discussion

### 5.1 New model with quadruple transition

With our double asymmetric percolation model, we observed a quadruple transition. As a result, when *T* is below the first epidemic threshold, there will only be microscopic non-outbreaks. When *T* is above the first epidemic threshold but below the second threshold, old homosexual males are most vulnerable to the outbreak. When *T* is above the second epidemic threshold but below the third threshold, both old heterosexual males and females become vulnerable to the outbreak. When *T* is above the third epidemic threshold but below the fourth threshold, young homosexual males become vulnerable to the outbreak. And when *T* is above the fourth epidemic threshold, young heterosexual males and females become vulnerable to the outbreak.

Based on our model, estimating the transmissibility and identifying vulnerable groups for better testing will provide better control and prevention over ZIKV outbreak.

### 5.2 Absence of epidemic thresholds in scale-free networks

The epidemic threshold, *θ*, in an uncorrelated network are shown to be

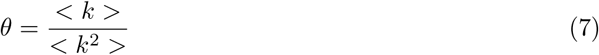

where *k* is the degree of nodes in a network [7].

For a scale-free network *P* (*k*) ∼ *k*^−*λ*^, with 2 < *λ* ≤ 3, the second moment < *k*^2^ > is diverging, i.e., < *k*^2^ > → + ∞. Therefore, the epidemic threshold *θ* is zero in a scale-free network with 2 < *λ* ≤ 3. This phenomena is then termed ‘null epidemic’ [6]. Similar results have been obtained in Erdős–Rényi model model when analyzing the critical point (where giant component emerges) [20].

Therefore, the disappearance of epidemic thresholds in a scale-free network in Section 4.3 can be explained by the null epidemic phenomena, meaning that outbreaks continue to exist for arbitrarily small transmissibility.

We note here that due to the fact that < *k*^2^ >→+∞ for a scale-free network with 2 < *λ* ≤ 3, some of the nodes in the networks can have a nearly infinite number of edges. It is highly likely for these nodes to start an outbreak even if the transmissibility is small. However, this setting is unlikely to be true for human sexual contact networks.

In addition, for a scale-free network with 3 < *λ*, epidemic thresholds reappear (see Figure 7). However, in our study, when 3 < *λ*, < *k* > is close to one. Such number of contacts is unrealistic for a human sexual contact network modeling.

We have also considered modeling scale-free networks with degree correlation. For example, individuals who have more sex partners are more likely to be connected with other individuals who have more sex partners. Nevertheless, [6] proved that *a scale-free degree distribution with exponent* 2 < *λ* ≤ 3 *is a sufficient condition for the absence of an epidemic threshold in unstructured networks with arbitrary two-point degree correlation function*. Therefore, scale-free networks with degree correlation will still not show any epidemic thresholds.

## 6 Conclusions and future work

In this project, we look at modeling of ZIKV transmission through sexual contacts. While gender plays an important role in modeling such a network, we argued that many other factors are also at play, including the two we examined: age and degree distribution. By integrating age as a second asymmetric factor, we observed a more complicated transition between outbreaks in different groups, described by 4 thresholds corresponding to different groups. The real-world scenario is even more complicated even with the absence of clustering by degrees. This means that outbreaks can happen at a smaller scale than previous predictions, and even with a low transmission rate. An epidemic can be sustained within a particular group, constantly spill to a larger population and may become a long standing concern.

Since a number of literature support the idea of modeling sexual activity as a scale-free network, we also tested the model on such type of network and find a transition at *λ* = 3. Before the transition, we see the “null-epidemic” scenario, which further supports the idea that the epidemic can be present for a long time in a small group even without the presence of mosquito vectors.

We believe our work can be further improved. On the one hand, we employ a simple separation between populations, using the age of 50 as a cutoff. Better classification is possible, for example, the younger group can be further split into children and adults, each has distinct properties. We also designed *w*_*i*_, *T*_*ij*_ and *α*_*i*|*j*_ for the new experiments, and these values are designed to largely follow the previous work with minimal changes, based on ball-park estimates. With the support of more real-world data, we shall have more freedom in designing the experiment, be able to build a better network in a more systematic way and observation can be closer to reality. We are also aware that there are not existing surveys for us to confirm or adjust our models due to a relatively low prevalence of the epidemic, and knowledge of real-world scenario will definitely help.

There are also potential improvements especially in the simulation part. Current algorithm requires knowing the exact composition of each giant components, which easily makes the algorithm run in *O*(*N* ^2^) time while time consumed on other steps only scales linearly with *N*. On the other hand, when the giant component become large, *T* is usually well past the last transition point and we are generally less interested in accuracy in this part. By preferential sampling over *T*, the scan parameter, we should be able to do a better simulation without sacrificing much into accuracy.

As the three mentioned metrics, *S, P* and < *s* > are important both theoretically and in practice, there is much more information to extract from both methods. For example, distribution of the size of small components might be of particular interest since it helps us to assess the feasibility of our “cutoff” and if the separation of giant and small components are clear for meaningful outcomes. Naturally, we can also look at < *p* >, from the composition of small out-components. While comparing *S* and *P* directly does not say much about the asymmetric properties of the graph, < *s* > and < *p* > have a better distinction seen from simulation, and we believe it’s worthy to analyze the difference of them if we want to be more informed about internal structure of the graph. And, in this study we only care about averages on these metrics, other measures, like standard deviation, can also be interesting in our analysis. For example, a high deviation on *P* or *S* might signal the outbreak is really unstable and a high deviation on < *s* > can be both from the intrinsic structure or from simulations. We can also expect that with different degree distributions comes to different statistics, as demonstrated before. In the future, better human sexual contact networks can be explored.

Finally, while we keep our focus on the non-vector transmission of ZIKV, asymmetric transmissions, or more generally asymmetric relations is commonly seen in many other diseases or even other interdisciplinary fields. We believe the ideas and methods presented here can be valuable to other studies in various fields.

## References

[1] Antoine Allard, Benjamin M Althouse, Samuel V Scarpino, and Laurent Hébert-Dufresne. Asymmetric percolation drives a double transition in sexual contact networks. Proceedings of the National Academy of Sciences, 114(34):8969–8973, 2017.

[2] Antoine Allard, Laurent Hébert-Dufresne, Jean-Gabriel Young, and Louis J Dubé. General and exact approach to percolation on random graphs. Physical Review E, 92(6):062807, 2015.

[3] Luis A Nunes Amaral, Antonio Scala, Marc Barthelemy, and H Eugene Stanley. Classes of small-world networks. Proceedings of the national academy of sciences, 97(21):11149–11152, 2000.

[4] Albert-László Barabási and Réka Albert. Emergence of scaling in random networks. science, 286(5439):509–512, 1999.

[5] Isaac I Bogoch, Oliver J Brady, Moritz UG Kraemer, Matthew German, Maria I Creatore, Shannon Brent, Alexander G Watts, Simon I Hay, Manisha A Kulkarni, John S Brownstein, et al. Potential for zika virus introduction and transmission in resource-limited countries in africa and the asia-pacific region: a modelling study. The Lancet infectious diseases, 16(11):1237–1245, 2016.

[6] Marián Boguná, Romualdo Pastor-Satorras, and Alessandro Vespignani. Absence of epidemic threshold in scale-free networks with degree correlations. Physical review letters, 90(2):028701, 2003.

[7] Stefan Bornholdt and Heinz Georg Schuster. Handbook of graphs and networks: from the genome to the internet. John Wiley & Sons, 2006.

[8] Gubio S Campos, Antonio C Bandeira, and Silvia I Sardi. Zika virus outbreak, bahia, brazil. Emerging infectious diseases, 21(10):1885, 2015.

[9] Van-Mai Cao-Lormeau, Claudine Roche, Anita Teissier, Emilie Robin, Anne-Laure Berry, Henri-Pierre Mallet, Amadou Alpha Sall, and Didier Musso. Zika virus, french polynesia, south pacific, 2013. Emerging infectious diseases, 20(6):1085, 2014.

[10] Christopher Chang, Kristina Ortiz, Aftab Ansari, and M Eric Gershwin. The zika outbreak of the 21st century. Journal of autoimmunity, 68:1–13, 2016.

[11] Mark R Duffy, Tai-Ho Chen, W Thane Hancock, Ann M Powers, Jacob L Kool, Robert S Lanciotti, Moses Pretrick, Maria Marfel, Stacey Holzbauer, Christine Dubray, et al. Zika virus outbreak on yap island, federated states of micronesia. New England Journal of Medicine, 360(24):2536–2543, 2009.

[12] G Ergün and GJ Rodgers. Growing random networks with fitness. Physica A: Statistical Mechanics and its Applications, 303(1):261–272, 2002.

[13] Güler Ergün. Human sexual contact network as a bipartite graph. Physica A: Statistical Mechanics and its Applications, 308(1):483–488, 2002.

[14] Brian D Foy, Kevin C Kobylinski, Joy L Chilson Foy, Bradley J Blitvich, Amelia Travassos da Rosa, Andrew D Haddow, Robert S Lanciotti, and Robert B Tesh. Probable non–vector-borne transmission of zika virus, colorado, usa. Emerging infectious diseases, 17(5):880, 2011.

[15] Daozhou Gao, Yijun Lou, Daihai He, Travis C Porco, Yang Kuang, Gerardo Chowell, and Shigui Ruan. Prevention and control of zika as a mosquito-borne and sexually transmitted disease: a mathematical modeling analysis. Scientific reports, 6:28070, 2016.

[16] Matt J Keeling and Pejman Rohani. Modeling infectious diseases in humans and animals. Princeton University Press, 2011.

[17] Adam J Kucharski, Sebastian Funk, Rosalind M Eggo, Henri-Pierre Mallet, W John Edmunds, and Eric J Nilles. Transmission dynamics of zika virus in island populations: a modelling analysis of the 2013–14 french polynesia outbreak. PLoS neglected tropical diseases, 10(5):e0004726, 2016.

[18] Fredrik Liljeros, Christofer R Edling, Luis A Nunes Amaral, H Eugene Stanley, and Yvonne Åberg. The web of human sexual contacts. Nature.

[19] Stacy Tessler Lindau and Natalia Gavrilova. Sex, health, and years of sexually active life gained due to good health: evidence from two us population based cross sectional surveys of ageing. BMj, 340:c810, 2010.

[20] Michael Molloy and Bruce Reed. A critical point for random graphs with a given degree sequence. Random structures & algorithms, 6(2-3):161–180, 1995.

[21] D Musso, EJ Nilles, and V-M Cao-Lormeau. Rapid spread of emerging zika virus in the pacific area. Clinical Microbiology and Infection, 20(10), 2014.

[22] Emanuele Nicastri, Concetta Castilletti, Giuseppina Liuzzi, Marco Iannetta, Maria R Capobianchi, and Giuseppe Ippolito. Persistent detection of zika virus rna in semen for six months after symptom onset in a traveller returning from haiti to italy, february 2016. Eurosurveillance, 21(32), 2016.

[23] Steven H Strogatz. Exploring complex networks. Nature, 410(6825):268–276, 2001.

[24] Benoit Visseaux, Emmanuel Mortier, Nadhira Houhou-Fidouh, Ségolène Brichler, Gilles Collin, Lucile Larrouy, Charlotte Charpentier, and Diane Descamps. Zika virus in the female genital tract. Lancet Infect Dis, 16(11):1220, 2016.

